# Towards the restoration of ancient hominid craniofacial anatomy: Chimpanzee morphology reveals covariation between craniometrics and facial soft tissue thickness

**DOI:** 10.1101/2021.01.08.425868

**Authors:** Ryan M Campbell, Gabriel Vinas, Maciej Henneberg

**Affiliations:** Adelaide Medical School, Biological Anthropology and Comparative Anatomy Research Unit, The University of Adelaide, Helen Mayo North, Floor 2, Room 24, Frome Road, Adelaide, South Australia,5000, Australia; Herberger Institute for Design and the Arts, Sculpture Department, Arizona State University, Art Building, 900 S Forest Mall, Tempe, Arizona, 85281, United States; Institute of Evolutionary Medicine, Faculty of Medicine, University of Zurich, Building 42, Floor G, Room 70, Winterthurerstr. 190, 8057, Zurich, Switzerland

**Keywords:** Evolution, Facial reconstruction, Forensic anthropology, Primate morphology.

## Abstract

In modern humans, facial soft tissue thicknesses have been shown to covary with craniometric dimensions. However, to date it has not been confirmed whether these relationships are shared with non-human apes. In this study, we analyze these relationships in chimpanzees (*Pan troglodytes*) with the aim of producing regression models for approximating facial soft tissue thicknesses in Plio-Pleistocene hominid individuals. Using CT scans of 19 subjects, 637 soft tissue, and 349 craniometric measurements, statistically significant multiple regression models were established for 26 points on the face and head. Examination of regression model validity resulted in minimal differences between observed and predicted soft tissue thickness values. Assessment of interspecies compatibility using a bonobo (*Pan paniscus*) and modern human *(Homo sapiens*) subject resulted in minimal differences for the bonobo but large differences for the modern human. These results clearly show that (1) soft tissue thicknesses covary with craniometric dimensions in *P. troglodytes*, (2) confirms that such covariation is uniformly present in both extant *Homo* and *Pan* species, and (3) suggests that chimp-derived regression models have interspecies compatibility with hominids who have similar craniometric dimensions to *P. troglodytes*. As the craniometric dimensions of early hominids, such as South African australopithecines, are more similar to *P. troglodytes* than those of *H. sapiens*, chimpanzee-derived regression models may be used for approximating their craniofacial anatomy. It is hoped that the results of the present study and the reference dataset for facial soft tissue thicknesses of chimpanzees it provides will encourage further research into this topic.

## Introduction

The primate family of Hominidae is comprised of the African apes, humans and all ancestors leading to these clades. Reconstructing soft tissue characters of extinct members of the Hominidae, called here hominids, has become an increasingly popular practice with many approximations of their faces presented in museum exhibitions, popular science publications and at conference presentations worldwide (1–3). In these contexts, reconstructions of the face and body have proven to be an effective vehicle for the dissemination of scientific information about human evolution. However, there is a recognized problem of variability among reconstructions of the same individual. A recent study comparing approximations of LB1, the holotype of *Homo floresiensis* (4, 5), reported that they vary significantly among one another (6). Similarly, in a systematic survey of 860 hominid reconstructions presented in 71 museums across Australia and Europe, it was found that inconsistencies are clearly prevalent in all other approximations of extinct hominid species (7). If practitioners were using reliable methods, this variability would not have occurred. Either the practitioners relied on unspecified sets of assumptions and/or the methods they used were unreliable. Furthermore, in its present state, the practice of hominid reconstructions is particularly vulnerable to attack from critics of human evolution who have used discrepancies between reconstructions of the same individual to undermine the reliability of evolutionary theory. Therefore, it is important to strengthen existing methods of reconstruction as much as possible to reduce this variability and avoid such criticisms.

The term used to describe the process of building a face over a skull varies in the literature between disciplines. In forensics, the name of the process most commonly referred to as ‘facial reconstruction’ was updated to ‘facial approximation’ because it is a more accurate description of the results, whereas in paleoanthropology the term ‘facial reconstruction’ is still being used (2). Regardless of what term is preferred, the results are always approximate and therefore we agree with previous authors and prefer the term ‘facial approximation’ (6, 8, 9). Scientific testing of facial approximation methods has been a major focus in craniofacial identification of human remains with research dating back at least to Welcker (10), with important contributions by Gerasimov (11, 12), Prag and Neave (13), and Wilkinson (14). Methods using means of soft tissues of the face have received the most attention (15–18), however, there is a recognized flaw in extrapolating means to individuals. As statistically robust as means may be, they only express means for specific populations. For reconstructing individuals, population means are not appropriate because they completely ignore variation among individuals. Regarding the approximation of extinct hominids, interspecies extrapolation of means derived from either modern humans or the extant great apes (*Gorilla*, *Pan*, and *Pongo*), as suggested in Hanebrink (17), is equally inappropriate because it also ignores variation among individuals.

One possible solution to this problem is to identify approximation methods that are compatible across all members of the Hominoidea superfamily. If a consistent pattern in covariation between soft tissue and craniometric measurements can be identified in extant hominids, then extinct hominids can reasonably be assumed to have followed suite. Such covariations were first explored in human material by Sutton (19) and extended in Simpson and Henneberg (20). Correlations were found and multiple linear regression models were used to generate equations for improving estimations of soft tissue thickness from craniometrics alone in modern humans, though this covariance has rarely been used in facial approximations. Reactions to the results of these studies are mixed. Stephan and Sievwright (21), using data measured with substantial random errors, report that regression models have low correlation coefficients that do not improve soft tissue thickness estimates above population means. However, Dinh, Ma (22) repeated the use of linear regression models and produced favorable results that encourage further exploration. Thus, for the purpose of hominid facial approximation hominids, the possibility of generating soft tissue thickness values that are individualized to a specific hominid specimen is undoubtably better than extrapolation of species-specific means.

The present study is motivated by the aforementioned concerns and while we hold that the findings reported here are valuable we raise three caveats at the outset: 1) As in previous studies of chimpanzee soft tissues (23–25), this study includes only a small sample of chimpanzees and therefore the conclusions from the results are subject to further testing on larger samples; 2) We also do not include other members of the African ape clade and so we cannot expand our findings to the entire Hominoidea superfamily; and 3) We do not claim to eliminate the need for informed speculation in hominid facial approximation entirely. Not all soft tissue characters of ancient hominids are addressed here, such as the facial features (eyes, nose, mouth, and ears) that arguably have a much greater impact on the variability between reconstructions of the same individual than soft tissue thicknesses alone. This work represents a step towards an empirical method that will strengthen the practice, but it is by no means a final solution to the problem.

The aims are: (1) To validate in chimpanzees (*Pan troglodytes*) that facial soft tissue thicknesses covary with craniometric dimensions, and (2) to produce soft tissue prediction models with interspecies compatibility from chimpanzee material that can be used in the facial approximation of extinct hominids.

## Materials and methods

Computed tomography scans of 28 chimpanzees were collected from two separate data repositories. Scans were accessed online, via the Digital Morphology Museum, KUPRI (dmm.pri.kyoto-u.ac.jp) and Morphosource (https://www.morphosouce.org), and obtained as Digital Imaging and Communications in Medicine (DICOM) format bitmap files. After excluding two neonates, one infant and six subjects showing obvious pathological effects or degradation caused by decomposition, the study sample contained 19 individuals of known age, sex and subject condition. The sex ratio was 1:1.71 (7 male and 12 female) and the mean age was 30.9 years (minimum = 9; maximum = 44; SD = 10.1). Subject condition was varied and included seven living, five fresh, five frozen, and two subjects that were preserved by immersion. Information on whether individual subjects were scanned in the supine or prone position was not available at the time of this study. Further information on whether subjects had been living in the wild prior to scanning was also not available but the study sample is assumed to consist of animals that were housed in captivity only. A complete list of all subjects used in this study is presented in the Supporting Information (S1 File).

Prior to measuring, all skulls were oriented in the Frankfurt horizontal plane determined according to its original definition by a horizontal line passing through the inferior border of the orbital rim (mid-infraorbital) and the top of the external auditory meatus (porion) on both sides of the skull. Facial soft tissue thickness was then measured at 39 cephalometric landmarks (Fig. 1; 17 medial and 22 bilateral) in OsiriX, v. 11.02 (Visage Imaging GmbH, Sand Diego, USA), which has been shown to produce accurate measurements that can be reliably compared between separate studies (26, 27). Cephalometric landmarks were selected based on common depths found in the facial approximation literature (20, 28–30). However, to allow for other aspects of the head beyond the face to be investigated, further points were added to include a wider range of points than normal, particularly points on the lateral areas of the head. The decision to include additional points can be explained as follows. The purpose of facial approximation of modern humans is to generate a specific recognition of a target individual, however, in facial approximation of ancient hominids the purpose is to show morphological differences between separate species. As morphological differences extend beyond the face to the rest of the head, the inclusion of further points allows for such comparisons to be made. A maximum total of 61 soft tissue depth measurements were possible per individual as well as 21 measurements of craniometric dimensions. All cephalometric points were positioned onto 3D volume renderings of the skulls prior to measuring with the exception of 6 points (gnathion, metopion, mid-mandibular body, mid-mandibular border, mid-nasal, mid-philtrum) that were more precisely aligned in 3D multiplanar reformatting by halving the inter-landmark distance between two adjacent points along the sagittal plane. When analyzing scans, it was noticed that some sutures of the cranium had been obliterated. However, as sutures are needed for the positioning of bregma, dacryon, ectoconchion, nasion, pterion, and zygomaxillare, these points were positioned and cross-checked against a reference skull of a chimpanzee from the Vernon-Roberts Anatomy and Pathology Museum, University of Adelaide. Depths at all landmarks were then measured perpendicular to the bone surface using the coronal, sagittal and transverse planes to control the direction of measurement. Xscope, v. 4.4.1 (ARTIS Software, Virginia, USA) was used to superimpose horizontal and vertical guides during the measurement procedure to keep measurements parallel to the reference planes. For thicknesses at bilateral landmarks, measurements were taken from both the right and the left side of the face and then the mean was calculated. Very few depths were unobtainable with the exception being those landmarks located in areas of incomplete DICOM data or where the soft tissues were outside of the anatomical position. For example, craniometrics crossing the occlusal line were not taken from subjects with open mandibles and soft tissue measurements were not taken for prosthion and/or infradentale in subjects with folded lips. Table 1 gives a complete list of all landmarks used in this study, their definitions, as well as their corresponding planes and angles of measurement. All soft tissue thickness data collected for chimpanzees has been made available by the authors on Figshare (https://figshare.com/s/8a3ba7df4dad9df7d70d).

**Fig 1.**
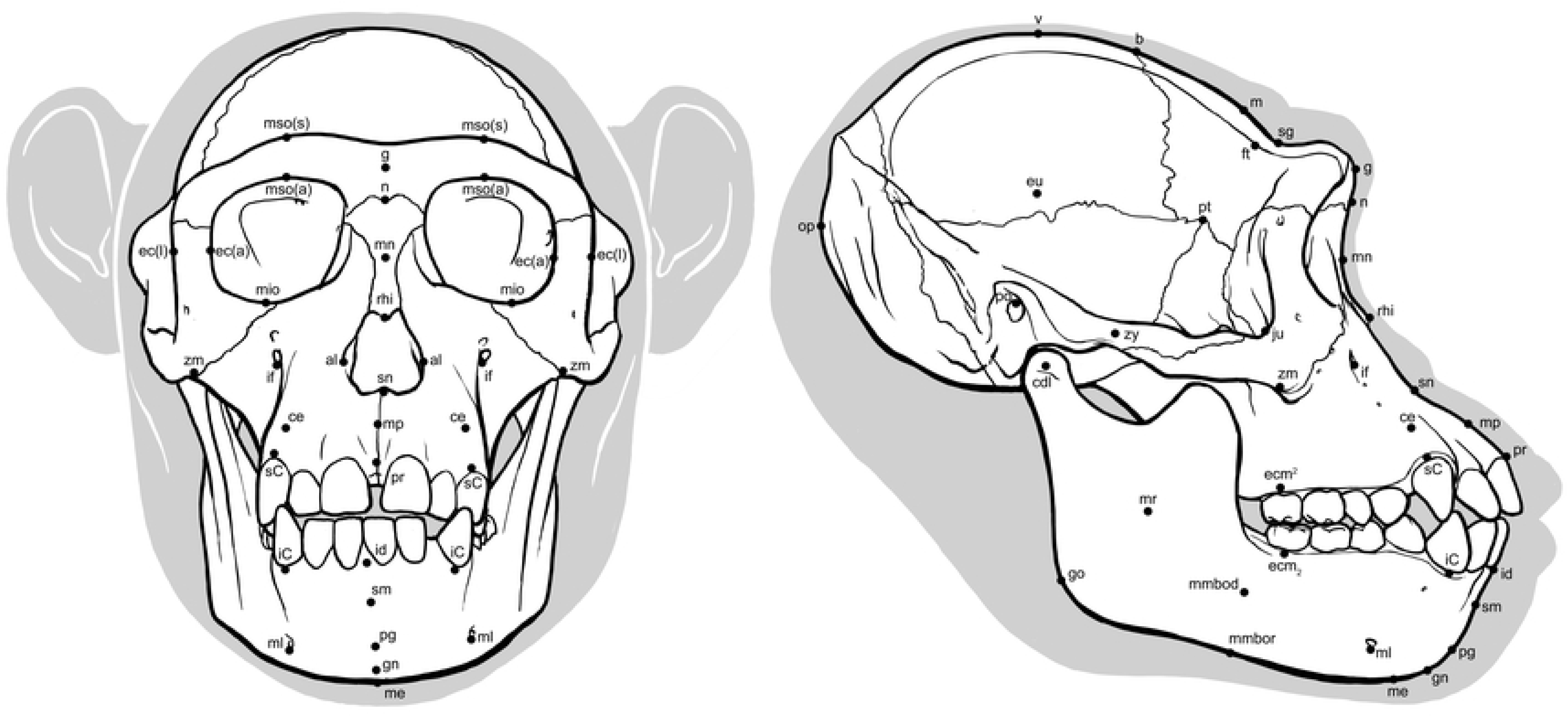
Locations of cephalometric landmarks on *Pan troglodytes* skull in norma frontalis and norma lateralis. See abbreviations in Table 1.

**Table 1.** Cephalometric landmarks including their abbreviations, definitions, planes of measurement, and angles of measurement. Points are listed in alphabetical order for ease of reference.

| **Landmark** | **Abbreviation** | **Definition** | **Plane of measurement** | **Angle of measurement** |
| --- | --- | --- | --- | --- |
| **Alare** | al | Instrumentally determined as the most lateral point on the nasal aperture in a transverse plane | transverse | parallel to reference plane |
| **Bregma** | b | Where the sagittal and coronal sutures meet | sagittal | perpendicular to bone |
| **Canine eminence** | ce | Most anterior point on the eminence of the maxillary canine | sagittal | perpendicular to bone |
| **Condylion laterale** | cdl | Most lateral point on the mandibular condyle | coronal | parallel to reference plane |
| **Ectoconchion**<br>**anteriorius** | ec(a) | Lateral point of the orbit at a line that bisects the orbit transversely | transverse | parallel to reference plane |
| **Ectoconchion**<br>**lateralis** | ec(l) | Most lateral point of the orbit, at a right angle to ec(l) in the transverse plane | transverse | parallel to reference plane |
| **Ectomolare<sup>2</sup>** | ecm<sup>2</sup> | Most lateral point on the buccal alveolar margin, at the center of the M<sup>2</sup> position | coronal | parallel to reference plane |
| **Ectomolare<sub>2</sub>** | ecm<sub>2</sub> | Most lateral point on the buccal alveolar margin, at the center of the M<sub>2</sub> position | coronal | parallel to reference plane |
| **Euryon** | eu | Instrumentally determined as the most lateral point of the cranial vault, on the parietal bone | transverse | parallel to reference plane |
| **Frontotemporale** | ft | Most anterior and medial point of the inferior temporal line, on the zygomatic process of the frontal bone | coronal | parallel to reference plane |
| **Glabella** | g | Most projecting anterior median point on lower edge of the frontal bone, on the brow ridge, in-between the superciliary arches and above the nasal root | sagittal | parallel to reference plane |
| **Gnathion** | gn | Median point halfway between pg and me | sagittal | perpendicular to bone |
| **Gonion** | go | Point on the rounded margin of the angle of the mandible, bisecting two lines. One following vertical margin of ramus and one following<br>horizontal margin of corpus of mandible | transverse | parallel to reference plane |
| **Infra canine** | iC | Point on the inferior alveolar ridge inferior to the<br>crown of the mandibular canine(s) | transverse | perpendicular to bone |
| **Infradentale** | id | Median point at the superior tip of the septum<br>between the mandibular central incisors | sagittal | parallel to reference<br>plane |
| **Infraorbital**<br>**foramen** | if | Most inferior point on the margin of the infraorbital<br>foramen | transverse | perpendicular to bone |
| **Jugale** | ju | Vertex of the posterior zygomatic angle, between<br>the vertical edge and horizontal part of the<br>zygomatic arch | transverse | parallel to reference<br>plane |
| **Mentale** | ml | Most inferior point on the margin of the mandibular<br>mental foramen | transverse | perpendicular to bone |
| **Menton** | me | Most inferior median point of the mental symphysis | sagittal | perpendicular to bone |
| **Metopion** | m | Median point, instrumentally determined on the frontal bone as the greatest elevation from a cord between n and b | sagittal | perpendicular to bone |
| **Mid-infraorbital** | mio | Point on the anterior aspect of the inferior orbital rim, at a line that vertically bisects the orbit | sagittal | parallel to reference plane |
| **Mid-mandibular body** | mmbod | Point on the lateral border of the corpus of the mandible midway between pg and go | coronal | perpendicular to bone |
| **Mid-mandibular border** | mmbor | Point on the inferior border of the corpus of the mandible midway between pg and go | coronal | perpendicular to bone |
| **Mid-nasal** | mn | Point on internasal suture midway between n and rhi | sagittal | perpendicular to bone |
| **Mid-philtrum** | mp | Median point midway between ss and pr | sagittal | perpendicular to bone |
| **Mid-ramus** | mr | Midpoint along the shortest antero-posterior depth of the ramus, in the masseteric fossa, and usually close to the level of the occlusal plane | coronal | perpendicular to bone |
| **Mid-supraorbital antierius** | mso(a) | Point on the anterior aspect of the superior orbital rim, at a line that vertically bisects the orbit | sagittal | parallel to reference plane |
| **Mid-supraorbital superius** | mso(s) | Most superior point on the brow ridge, at a right angle to mso(a) in the sagittal plane | sagittal | parallel to reference plane |
| **Nasion** | n | Intersection of the nasofrontal sutures in the medial plane | sagittal | parallel to reference plane |
| **Opisthocranion** | op | Most posterior median point on the occipital bone, instrumentally determined as the greatest chord length from g. Note that this points is not necessarily at the tip of the nasal spine. | sagittal | parallel to reference plane |
| **Pogonion** | pg | Most anterior median point on the mental eminence of the mandible | sagittal | perpendicular to bone |
| **Porion** | po | Most superior point on the upper margin of the external auditory meatus | sagittal | parallel to reference plane |
| **Prosthion** | pr | Median point between the central incisors on the anterior most margin of the maxillary alveolar rim | sagittal | parallel to reference plane |
| **Pterion** | pt | A circular region, marked by the sphenoparietalis suture at its center. This region marks the thinnest part of the cranial vault | transverse | parallel to reference plane |
| **Rhinion** | rhi | Most rostral (end) point on the internasal suture. It cannot be determined accurately if nasal bones are broken distally | sagittal | perpendicular to bone |
| **Subnasale** | sn | The deepest point seen in the profile view below the anterior nasal spine | sagittal | perpendicular to bone |
| **Supra canine** | sC | Point on superior alveolar ridge superior to the crown of the maxillary canine(s) | transverse | perpendicular to bone |
| **Supraglabellare** | sg | Deepest part of the supraglabella fossa in the median plane | sagittal | parallel to reference plane |
| **Supramentale** | sm | Deepest median point in the groove superior to the mental eminence | sagittal | perpendicular to bone |
| **Vertex** | v | Most superior point of the skull | sagittal | parallel to reference plane |
| **Zygion** | zy | Instrumentally determined as the most lateral point on the zygomatic arch | coronal | parallel to reference plane |
| **Zygomaxillare** | zm | Most inferior point on the zygomaticomaxillary suture | transverse | parallel to reference plane |

192 Landmarks are positioned and measurements taken assuming the FH position.

Intra-observer soft tissue and craniometric measurement reliability was assessed by test-retest measurements following recently discussed data collection protocols in Stephan et al. (31). Measurements were taken on a subsample of five individuals that were randomly selected by a volunteer. Specimens were PRI-9783, PRI-Akira, PRI-10814, PRI-Mari and PRI-Reiko. Measurements of each specimen were conducted on separate days over a four-hour period with retest measurements taken seven days after initial assessment for 40 total measuring hours (96 hours including the entire sample). Intra-observer measurement reliability was calculated using the technical error of measurement (TEM) equation:

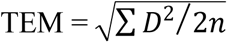

Where D is the difference between the values obtained for measurements taken on two separate occasions for each variable raised to the second power and *n* is the number of individuals measured. The relative TEM (r-TEM) was also calculated by dividing TEM by the mean measurement to convert to a percentage value.

Two-tail t-tests were used to analyze differences in mean soft tissue thicknesses and craniometric dimensions between males and females as well as between living and deceased subjects. Significance levels were set at *p* < 0.05 but altered according to the Bonferroni adjustment for 39 comparisons of soft tissue thicknesses and 21 comparisons of craniometric dimensions. Post-hoc power analyses were also performed to validate conclusions from two-tail t-tests. These analyses were conducted in Microsoft^®^ Excel^®^, v. 16.39 for Mac.

Data distribution was checked using the Kolmogorov-Smirnov and Shapiro-Wilk tests. All variables’ distributions did not differ significantly from the normal distribution, so no attenuation correction was made. Stepwise multivariate linear regression analysis was then performed to examine the relationships between facial soft tissue thicknesses and craniometric dimensions. Due to the small sample size, missing values in the data set were imputed with variable means rather than using list-wise or pair-wise deletion. These analyses were carried out with the Statistical Package for the Social Sciences (SPSS^®^) software, v. 26.0 for Mac (SPSS Inc, Chicago, II, USA).

To assess the validity of the regression models, they were tested on an in-group sample of 19 subjects. Craniometric measurements for each subject, taken from the results of the above described measurement procedure, were employed with the appropriate regression models to predict facial soft tissue thickness at 26 landmarks (8 medial and 18 bilateral). Empirical comparisons were then calculated as the simple difference, z-score and relative percent difference between predicted and observed soft tissue thicknesses. These analyses were conducted in Microsoft^®^ Excel^®^, v. 16.39 for Mac.

To validate the interspecies compatibility of the regression models, an out-of-group test was conducted on computed tomography scans of one living modern human male of European decent and one wet specimen of a deceased male bonobo *(Pan paniscus*; subject S9655). Bonobos and chimpanzees are highly similar to each other in many respects, however, they are classified as distinct species (32). S9655 is a sub-adult (4 years) male and thus outside the age-range of this study sample. Therefore, the bonobo served as an independent test on a sub-adult individual belonging to a separate species that was not used to generate the regression models. Moreover, to the knowledge of the authors, this is the only bonobo scan that is publicly available and as such removes the possibility of selection bias. The bonobo was accessed online via Morphosource (https://www.morphosouce.org). The human subject was donated specifically for the purpose of this study to the University of Adelaide by an anonymous donor. Both subjects were obtained as DICOM format bitmap files. Soft tissue measurements used in the comparison between observed and predicted values were taken following the aforementioned protocol.

To demonstrate the practical utility of the regression models, a 3D facial approximation was performed on the skull of subjects PRI-Cleo, S9655, as well as on an *Australopithecus africanus* skull (a composite reconstruction of specimens Sts 5 and Sts 52) previously described in Strait et al. (33) and Benazzi et al. (34, 35). The Sts 5 specimen, dated to 2.14 Ma (36) and found in 1947 at the South African Sterkfontein site by Robert Broom and John Robinson (37), is a perfect candidate for demonstrating the utility of the facial approximation method. Despite the edentulous maxilla and a break in the cranium associated with a dynamite explosion at the time of discovery, Sts 5 is an exceptionally preserved specimen relative to most other Plio-Pleistocene hominid skulls (38–40). It is worth nothing here that while the identification of Sts 5 as a female has been and is still the subject of ongoing debate (41–43), the cranial features of the Sts 5 specimen suggest that it was certainly not a subadult individual; therefore, it is well within the age-range of this study sample (44, 45). The Sts 52 mandible was reconstructed using state-of-the-art digital methods from research quality casts of the original specimen. For a full description of the digital reconstruction process, see Supporting Information in Benazzi et al. (34).

To begin the facial approximation procedure, soft tissue thicknesses were estimated for all three subjects by taking craniometric measurements from the skulls in digital format and inserting these measurements into the appropriate regression models. The resulting values were then used to design and place pegs corresponding to the results onto the skulls at their appropriate cephalometric points in Autodesk Maya, 2018 (Autodesk, San Rafael, CA). Both skulls were then 3D printed as recommended by Walker and Humphries (46). Each skull was printed separately with articulated mandibles on the M200 3D printer (Zortrax^®^) in acrylonitrile butadiene styrene via fused deposition modelling. Post-processing of prints involved the removal of all support material and then mounting the skulls in the Frankfurt horizontal plane. As a precautionary measure, all three of the 3D printed skulls were measured and cross-checked against measurements taken from their digital counterparts. No discrepancies were observed. The soft tissues were then constructed using an oil-based modelling medium by GV using the pegs to guide the thicknesses at each cephalometric point. Given that facial features (eyes, nose, mouth and ears) were not the focus of the present study and are likely to be based on intuition rather than empirical science, especially in the case of *A. africanus,* the eyes were closed, the nose and mouth were left undefined and the ears were omitted from all three approximations.

## Results

The descriptive statistics for soft tissue measurements and the craniometrics are presented in Tables 2 and 3 respectively, along with the intra-observer TEM and r-TEM for each variable. The mean intra-observer r-TEM for measurements of soft tissue thickness was 2.38%. The lowest intra-observer r-TEM was observed for prosthion (0.45%) and the largest was rhinion (7.00%). The mean intra-observer r-TEM for measurements of soft tissue thickness is lower than the mean intra-observer r-TEM of 8% recorded by Stephan and Sievwright (21), which involved measurements of living subjects by B-mode ultrasound. For the craniometrics, the mean intra-observer r-TEM was 0.25%. The lowest intra-observer r-TEM was observed for the distance from vertex to subnasale (0.05%) and the largest was for bigonial breadth (0.93%). The mean intra-observer r-TEM for the craniometrics also stands in contrast to the mean intra-observer r-TEM of 2% recorded by Stephan and Sievwright (21). These results follow the usual trend whereby soft tissue measurements are shown to pose a greater challenge to measurement accuracy than the craniometrics. However, they also show that measurements taken from CT scans in OsiriX in the present study appear to be more accurate than measurements taken in previous studies via ultrasound (21). It may be suggested that the TEM values reported here are underestimates because the repeat measurements were performed on the same scans rather than replicating the whole measurement process to include the reacquisition of scans. However, not only are CT scans of great apes exceptionally rare but also reacquisition of scans of living apes would expose these endangered animals to needless levels of ionizing radiation and is, therefore, unethical.

**Table 2.** Descriptive statistics of facial soft tissue thickness in mm for *Pan troglodytes*.

| **Variable<sup>a</sup>** | **Mean** | **SD** | **Minimum** | **Maximum** | ***n*** | **TEM<sup>b</sup>**<br>**(mm)** | **rTEM<sup>c</sup>**<br>**(%)** |
| --- | --- | --- | --- | --- | --- | --- | --- |
| **Median points** |  |  |  |  |  |  |  |
| **v** | 5.36 | 2.28 | 2.40 | 8.83 | 13 | 0.05 | 0.97 |
| **b** | 4.17 | 1.50 | 2.23 | 7.89 | 13 | 0.18 | 4.55 |
| **m** | 3.29 | 1.04 | 1.79 | 5.44 | 12 | 0.19 | 5.88 |
| **sg** | 3.80 | 1.56 | 2.00 | 8.49 | 15 | 0.08 | 2.18 |
| **g** | 4.00 | 1.09 | 2.39 | 6.13 | 17 | 0.20 | 6.45 |
| **n** | 3.24 | 0.83 | 1.98 | 5.27 | 19 | 0.06 | 1.99 |
| **mn** | 2.33 | 0.50 | 1.59 | 3.42 | 19 | 0.06 | 2.39 |
| **rhi** | 4.07 | 1.53 | 1.00 | 8.31 | 18 | 0.26 | 7.00 |
| **sn** | 7.33 | 1.94 | 3.72 | 12.50 | 18 | 0.15 | 1.70 |
| **mp** | 12.29 | 4.49 | 6.42 | 20.20 | 12 | 0.51 | 3.71 |
| **pr** | 14.06 | 2.16 | 10.60 | 17.85 | 9 | 0.07 | 0.45 |
| **id** | 15.54 | 3.71 | 10.10 | 22.55 | 14 | 0.18 | 0.94 |
| **sm** | 17.79 | 3.98 | 10.60 | 24.35 | 17 | 0.43 | 2.21 |
| **pg** | 12.37 | 4.18 | 4.80 | 19.00 | 16 | 0.55 | 3.35 |
| **gn** | 8.01 | 3.31 | 3.17 | 13.70 | 17 | 0.29 | 2.45 |
| **me** | 5.86 | 2.04 | 3.10 | 10.70 | 17 | 0.15 | 2.35 |
| **Bilateral points** |  |  |  |  |  |  |  |
| **mso(a)** | 7.25 | 1.63 | 4.27 | 9.73 | 15 | 0.11 | 1.80 |
| **mso(s)** | 4.69 | 1.39 | 2.83 | 7.30 | 15 | 0.13 | 2.34 |
| **mio** | 5.86 | 2.25 | 3.01 | 12.35 | 19 | 0.11 | 1.94 |
| **ec(a)** | 6.01 | 1.84 | 1.91 | 8.81 | 18 | 0.21 | 2.98 |
| **ec(l)** | 7.14 | 2.91 | 2.43 | 12.60 | 17 | 0.07 | 0.81 |
| **zy** | 7.13 | 2.36 | 2.54 | 11.70 | 18 | 0.21 | 2.70 |
| **cdl** | 20.83 | 5.72 | 10.33 | 32.95 | 18 | 0.20 | 0.89 |
| **ft** | 26.97 | 9.25 | 13.20 | 41.73 | 13 | 0.53 | 1.67 |
| **pt** | 34.64 | 9.10 | 19.75 | 56.75 | 15 | 0.28 | 0.77 |
| **eu** | 20.16 | 7.45 | 10.21 | 36.30 | 12 | 0.23 | 1.15 |
| **ju** | 7.99 | 2.96 | 3.15 | 14.60 | 18 | 0.16 | 1.74 |
| **if** | 9.52 | 1.84 | 5.70 | 12.48 | 19 | 0.07 | 0.77 |
| **zm** | 9.17 | 2.66 | 2.67 | 13.23 | 18 | 0.09 | 1.03 |
| **ce** | 11.21 | 3.74 | 6.05 | 18.25 | 18 | 0.26 | 2.27 |
| **ecm<sup>2</sup>** | 22.50 | 6.52 | 14.93 | 37.10 | 19 | 0.60 | 2.88 |
| **ecm<sub>2</sub>** | 20.04 | 5.98 | 12.80 | 33.50 | 19 | 0.60 | 3.17 |
| **sC** | 9.88 | 2.19 | 6.30 | 13.08 | 14 | 0.20 | 2.02 |
| **iC** | 11.19 | 2.21 | 8.41 | 16.05 | 18 | 0.13 | 1.04 |
| **ml** | 7.17 | 2.14 | 3.54 | 12.25 | 17 | 0.31 | 5.32 |
| **mr** | 26.63 | 6.70 | 14.35 | 44.70 | 18 | 0.17 | 0.59 |
| **go** | 11.44 | 3.64 | 5.22 | 19.78 | 18 | 0.51 | 3.35 |
| **mmborder** | 5.24 | 2.42 | 2.15 | 10.55 | 17 | 0.07 | 1.35 |
| **mmbody** | 13.95 | 7.15 | 4.27 | 34.05 | 18 | 0.27 | 1.58 |

<sup>a</sup> See variable abbreviations in Table 1. <sup>b</sup> Technical error of measurement. <sup>c</sup> Relative technical error of measurement.

**Table 3.** Descriptive statistics of craniometric dimensions in mm for *Pan troglodytes*.

| Variable<sup>a</sup> | Measurement | Mean | SD | Minimum | Maximum | *n* | TEM<sup>b</sup> (mm) | rTEM<sup>c</sup> (%) |
| --- | --- | --- | --- | --- | --- | --- | --- | --- |
| **v-po** | auricular height | 65.61 | 4.14 | 58.80 | 73.40 | 13 | 0.10 | 0.15 |
| **v-g** | distance from vertex to glabella | 89.63 | 6.02 | 74.60 | 99.00 | 12 | 0.07 | 0.08 |
| **v-n** | distance from vertex to nasion | 92.45 | 5.56 | 78.10 | 100.00 | 13 | 0.11 | 0.12 |
| **v-sn** | distance from vertex to subnasale | 135.25 | 7.39 | 118.30 | 149.50 | 13 | 0.07 | 0.05 |
| **v-gn** | distance from vertex to gnathion | 198.87 | 11.98 | 178.80 | 228.10 | 12 | 0.45 | 0.23 |
| **g-n** | supraorbital torus height | 11.03 | 2.51 | 6.09 | 15.85 | 18 | 0.06 | 0.56 |
| **g-sn** | distance from glabella to subnasale | 60.84 | 7.26 | 36.60 | 68.20 | 18 | 0.13 | 0.21 |
| **g-gn** | total face height | 140.92 | 11.63 | 125.20 | 167.70 | 17 | 0.22 | 0.16 |
| **g-op** | maximum head length | 142.51 | 7.29 | 132.50 | 153.60 | 15 | 0.19 | 0.13 |
| **n-sn** | nasal height | 53.43 | 4.55 | 44.30 | 61.30 | 19 | 0.15 | 0.29 |
| **n-pr** | upper face height | 86.63 | 7.91 | 75.10 | 101.10 | 19 | 0.11 | 0.13 |
| **id-gn** | mandibular symphysis height | 41.44 | 5.69 | 31.00 | 51.70 | 19 | 0.11 | 0.27 |
| **eu-eu** | maximum head breadth | 97.98 | 3.90 | 90.95 | 104.00 | 13 | 0.13 | 0.13 |
| **ft-ft** | minimum frontal breadth | 45.51 | 9.21 | 32.00 | 66.40 | 15 | 0.08 | 0.18 |
| **zy-zy** | bizygomatic breadth | 131.11 | 11.05 | 116.50 | 156.40 | 19 | 0.07 | 0.05 |
| **c-c** | intercanine breadth | 63.00 | 7.38 | 47.60 | 75.10 | 19 | 0.39 | 0.62 |
| **go-go** | bigonial breadth | 96.39 | 10.17 | 79.50 | 118.50 | 19 | 0.90 | 0.93 |
| **al-al** | nasal breadth | 27.95 | 1.90 | 24.40 | 31.15 | 19 | 0.04 | 0.16 |
| **ec-ec** | biorbital breadth | 91.20 | 5.26 | 84.00 | 102.40 | 19 | 0.35 | 0.39 |
| **obh** | orbital height | 34.54 | 2.16 | 29.90 | 37.50 | 19 | 0.08 | 0.23 |
| **obb** | orbital breadth | 36.13 | 2.32 | 31.60 | 41.38 | 19 | 0.08 | 0.23 |

<sup>a</sup> See variable abbreviations in Table 1.
<sup>b</sup> Technical error of measurement.
<sup>c</sup> Relative technical error of measurement.

Two-tail t-tests show that there were no significant differences between living and deceased chimpanzee means across all 39 soft tissue thicknesses and 21 craniometric dimensions. This implies that the use of fresh, frozen and immersed subjects in this study did not compromise the validity of the results. However, post-hoc power analysis revealed that the probability of detecting a significant difference was low and that these results should be taken as tentative. In contrast, post-hoc power analysis revealed that comparisons of sexual dimorphisms were valid. Our results are in agreement with prior research in that there are no significant differences between male and female means (17), with the exception of two soft tissue landmarks (vertex and frontotemporale) and two craniometrics (bizygomatic breadth and inter-canine breadth). The greatest difference between male and female soft tissue thicknesses was observed for vertex and the lowest was observed for bizygomatic breadth. Note that this means there were only two instances out of 39 soft tissue depths that were not significantly different between the sexes.

In stepwise multivariate linear regression analyses, statistically significant (*p* ≤ 0.05) correlations between soft tissue depth measurements and craniometric dimensions were found. Of the 39 cephalometric landmarks assessed, statistically significant regression models were established for 26 landmarks (8 medial and 18 bilateral) and these are given in Table 4. Scatterplots showing four examples of bivariate relationships are shown in Fig 2. As such, it is now possible to reconstruct soft tissue thickness at 44 individual points on the face of chimpanzees using regression models alone. The mean standard error of the estimate (SEE) was 2.39 mm, ranging from 0.42 mm for menton to 5.29 mm for ectomolare_2_, and the mean multiple correlation coefficients (R = 0.67) far exceed those produced in previous studies of human material (20, 21). The model with the highest correlation coefficient was pterion (r = 0.93) and the lowest was mid-nasal (r = 0.46). A factor that should be considered here is that if this study sample of chimpanzees was indeed composed of subjects that were scanned in both the supine and prone positions, then the correlations observed here are likely underestimates and the true strengths of correlations higher than those reported.

**Fig 2.**
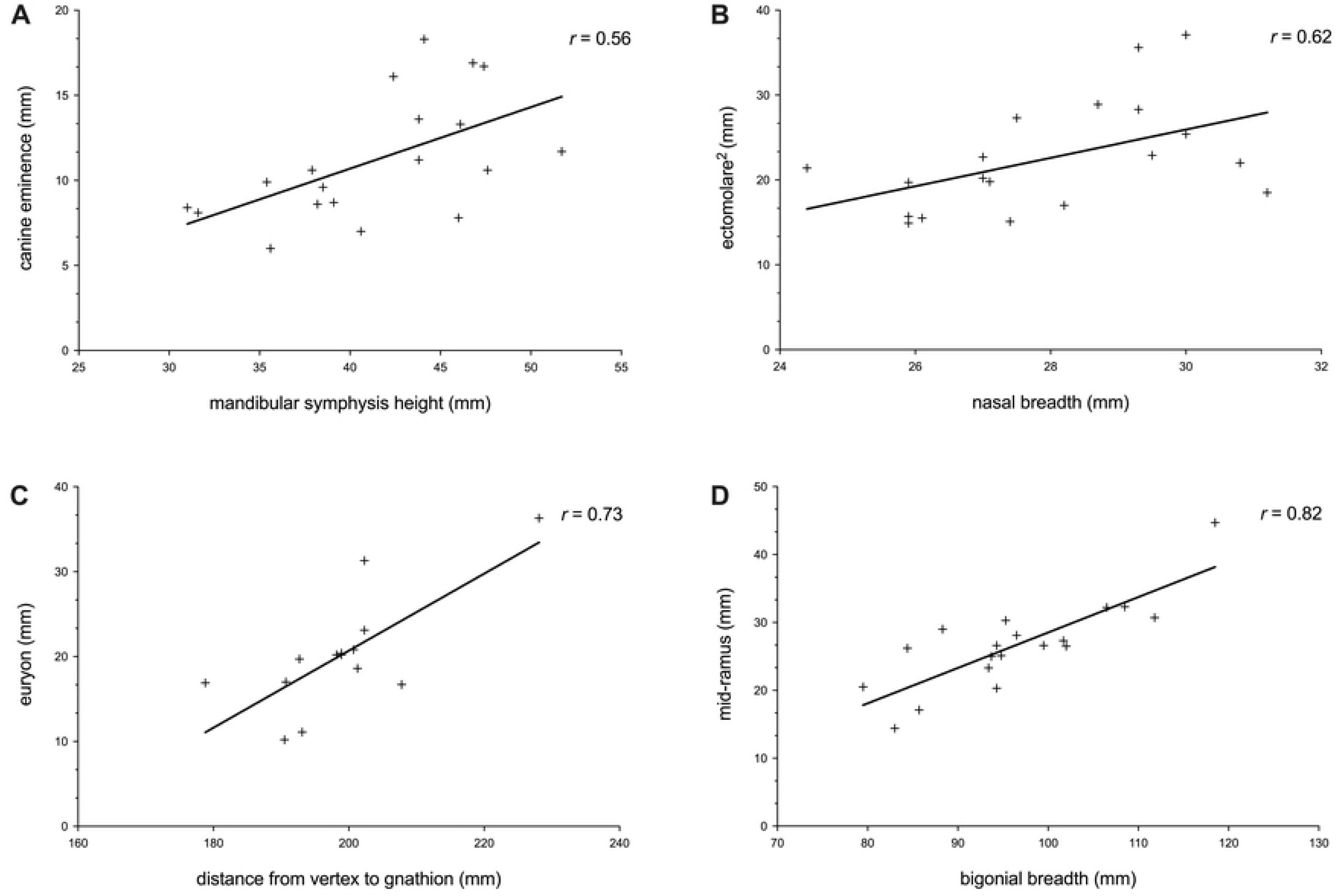
Scatterplots showing four examples of covariation between facial soft tissue and craniometric measurements in chimpanzees. (A) Correlation between canine eminence and mandibular symphysis height. (B) Correlation between ectomolare^2^ and nasal breadth. (C) Correlation between euryon and distance from vertex to gnathion. (D) Correlation between mid-ramus and bigonial breadth.

**Table 4.** Chimpanzee-derived linear regression models for medial and bilateral facial soft tissue thicknesses.

| **Variable** | **Linear regression model<sup>a</sup>** | **R** | **SE** | **P** |
| --- | --- | --- | --- | --- |
| **Median points** |  |  |  |  |
| **Vertex** | $0.095 (zy-zy) - 0.310 (g-n) - 3.680$ | 0.71 | 1.40 | 0.04 |
| **Bregma** | $0.052 (v-gn) - 0.205 (g-n) + 0.048 (g-gn) - 10.689$ | 0.87 | 0.70 | 0.04 |
| **Metopion** | $0.113 (v-p) - 0.158 (g-n) + 0.055 (v-sn) - 9.832$ | 0.88 | 0.42 | 0.01 |
| **Supraglabella** | $0.149 (v-sn) - 16.370$ | 0.66 | 1.07 | <0.01 |
| **Mid-nasal** | $-0.040 (id-gn) + 3.996$ | 0.46 | 0.46 | 0.05 |
| **Subnasale** | $0.510 (al-al) - 6.942$ | 0.52 | 1.66 | 0.02 |
| **Prosthion** | $0.169 (v-g) - 0.064 (g-gn) + 7.897$ | 0.68 | 1.12 | 0.02 |
| **Menton** | $0.131 (c-c) - 0.187 (v-n) + 14.818$ | 0.73 | 1.40 | 0.02 |
| **Bilateral points** |  |  |  |  |
| **Mid-supraorbital superius** | $-0.250 (g-n) + 7.449$ | 0.50 | 1.09 | 0.03 |
| **Ectoconchion lateralis** | $0.425 (v-p) + 0.127 (go-go) - 32.987$ | 0.81 | 1.71 | 0.01 |
| **Zygion** | $-0.655 (g-n) + 14.354$ | 0.70 | 1.70 | <0.01 |
| **Condylion laterale** | 1.144 (v-p) – 54.201 | 0.69 | 4.11 | <0.01 |
| **Frontotemporale** | 0.358 (n-pr) + 0.361 (go-go) – 0.277 (ft-ft) – 0.934 (g-n) – 15.905 | 0.90 | 3.72 | 0.03 |
| **Pterion** | 0.366 (zy-zy) + 0.691 (n-sn) + 0.965 (v-p) – 113.639 | 0.93 | 3.26 | <0.01 |
| **Euryon** | 0.454 (v-gn) – 70.061 | 0.73 | 4.10 | <0.01 |
| **Jugale** | 0.573 (v-p) – 29.626 | 0.67 | 2.18 | <0.01 |
| **Infraorbital foramen** | 0.150 (id-gn) + 3.296 | 0.47 | 1.67 | 0.04 |
| **Zygomaxillare** | 0.416 (v-p) – 18.139 | 0.54 | 2.23 | 0.02 |
| **Canine eminence** | 0.360 (id-gn) – 3.712 | 0.56 | 3.09 | 0.01 |
| **Ectomolare<sup>2</sup>** | 0.369 (zy-zy) – 25.860 | 0.62 | 5.24 | <0.01 |
| **Ectomolare<sub>2</sub>** | 1.601 (al-al) – 24.708 | 0.51 | 5.29 | 0.03 |
| **Infra canine** | – 0.110 (v-gn) + 33.154 | 0.48 | 1.93 | 0.04 |
| **Mid-ramus** | 0.523 (go-go) – 23.785 | 0.82 | 3.88 | <0.01 |
| **Gonion** | 0.628 (v-p) – 29.791 | 0.60 | 2.91 | 0.01 |
| **Mid-mandibular border** | 0.316 (v-p) – 15.491 | 0.47 | 2.07 | 0.04 |

<sup>a</sup> See variable abbreviations in Table 1.

The results of the regression models applied to the in-group sample of 19 subjects are shown in Table 5. Overall, the performance of the regression models was accurate as the differences between observed and predicted soft tissue thickness values were small (mean difference in mm = 1.96 mm; mean z-score = 0.54; mean relative difference 18.13%). The minimum difference in mm was observed in the regression model for *mid-nasal* (0.30 mm) and the maximum for ectomolare ^2^ (5.18 mm) The minimum z-score was observed in the regression model for pterion (0.28) and the maximum for mid-supraobital superius (0.96). The minimum relative difference was observed in the regression model for prosthion (7.62%) and the maximum for mid-mandibular border (28.50%).

**Table 5.** Average differences between predicted soft tissue thicknesses and ground truth values in this study sample of chimpanzees (*Pan troglodytes*).

| Variable<sup>a</sup> | Difference (mm) | Z-score | Relative difference (%) | *n* |
| --- | --- | --- | --- | --- |
| **Median points** |  |  |  |  |
| **v** | 1.22 | 0.51 | 25.33 | 11 |
| **b** | 0.57 | 0.35 | 15.06 | 11 |
| **m** | 0.33 | 0.31 | 10.59 | 10 |
| **sg** | 0.96 | 0.56 | 24.02 | 11 |
| **mn** | 0.30 | 0.61 | 11.62 | 11 |
| **sn** | 1.55 | 0.70 | 19.18 | 10 |
| **pr** | 1.14 | 0.42 | 7.62 | 5 |
| **me** | 1.23 | 0.52 | 20.13 | 10 |
| **Bilateral**<br>**points** |  |  |  |  |
| **mso(s)** | 1.45 | 0.96 | 26.64 | 11 |
| **ec(l)** | 1.18 | 0.38 | 18.35 | 11 |
| **zy** | 1.44 | 0.54 | 21.02 | 11 |
| **cdl** | 3.32 | 0.47 | 16.69 | 11 |
| **ft** | 2.78 | 0.30 | 9.85 | 10 |
| **pt** | 2.82 | 0.28 | 8.77 | 10 |
| **eu** | 3.79 | 0.46 | 21.32 | 10 |
| **ju** | 1.83 | 0.56 | 23.35 | 11 |
| **if** | 1.42 | 0.78 | 14.82 | 11 |
| **zm** | 2.03 | 0.63 | 26.00 | 10 |
| **ce** | 1.87 | 0.55 | 16.55 | 10 |
| **ecm<sup>2</sup>** | 5.18 | 0.79 | 22.78 | 11 |
| **ecm<sub>2</sub>** | 3.35 | 0.62 | 15.99 | 11 |
| **iC** | 1.60 | 0.60 | 13.90 | 10 |
| **mr** | 3.33 | 0.54 | 14.45 | 11 |
| **go** | 2.57 | 0.59 | 21.64 | 11 |
| **mmbor** | 1.56 | 0.61 | 28.50 | 10 |
| **mmbod** | 2.27 | 0.38 | 17.11 | 11 |

351 <sup>a</sup> See variable abbreviations in Table 1.

Fig 3 shows the results of the regression models applied in 3D facial approximations of subject PRI-Cleo (Fig. 3A), the out-of-group bonobo subject S9655 (Fig. 3B), and the composite skull of *A. africanus* (hereafter referred to as Sts 5; Fig. 3C). The differences between observed and predicted soft tissue thicknesses for PRI-Cleo and S9655 were small (Fig 4). The mean difference across 25 landmarks was 1.3 mm for PRI-Cleo (minimum = 0.1 mm; maximum = 5.8 mm) and 1.5mm for S9655 (minimum = 0 mm; maximum = 4.1 mm), which demonstrates that chimpanzee-derived regression models have closely predicted the soft tissue thicknesses for a sub-adult bonobo. In contrast, in the human subject the mean difference across 26 landmarks was 14.5 mm (minimum = 0.1 mm; maximum = 61.5 mm). This difference is much higher than those reported for PRI-Cleo and S9655 and in one case (menton) the regression model produced a negative result, which clearly indicates a fundamental problem with chimp-derived regression models for predicting modern human soft tissue thicknesses. As it is not possible for tissues thicknesses to be negative or equal to zero a 3D facial approximation of the human subject was not produced.

**Fig 3.**
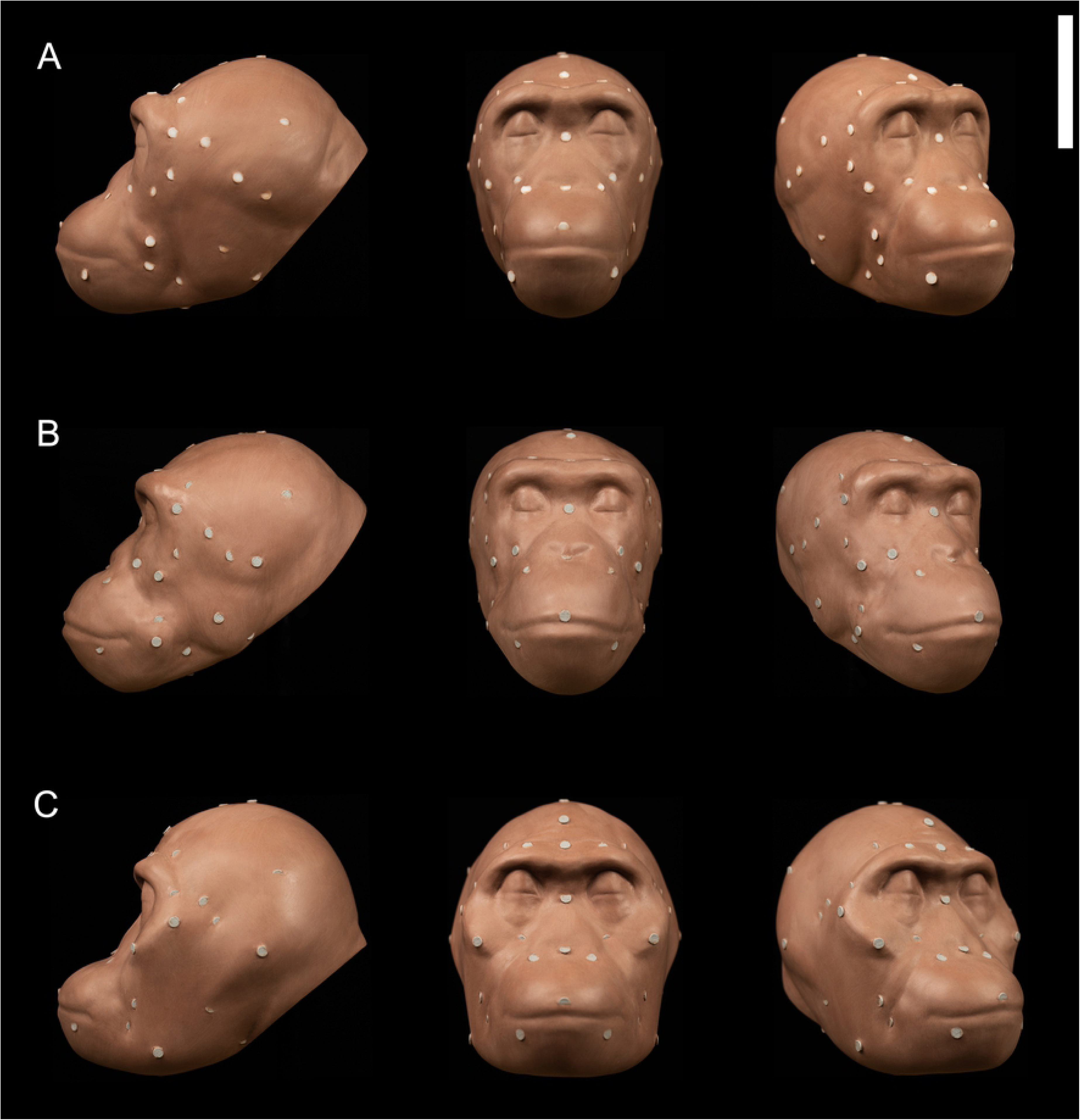
3D facial approximations of PRI-Cleo (*Pan troglodytes*; A), 29655 (*Pan paniscus*; B), and composite skull of *Australopithecus africanus* (Sts 5 and Sts 52; C) in right three quarter view (30° rotation from full face), norma frontalis and, norma lateralis. Note that the angle of the head in each facial approximation follows standard orientation methods established for modern humans. For *P. troglodytes*, *P.* paniscus, and *A. africanus* this angle may be unjustified biomechanically (Johanson, 1981). Scale bar = 10 cm.

**Fig 4.**
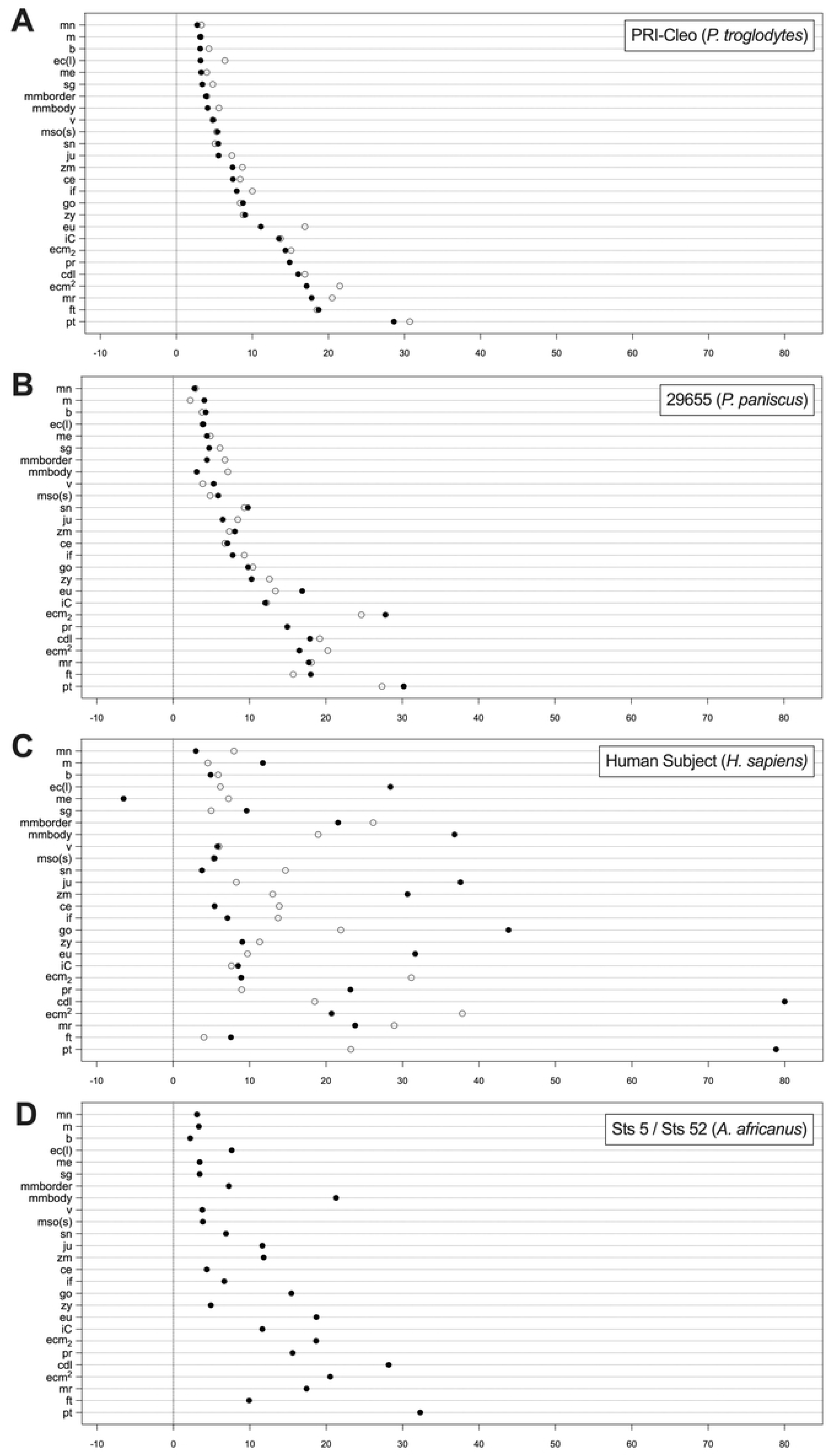
Depth chart comparison of observed (○) and predicted (●) facial soft tissue thickness values between facial approximations of PRI-Cleo (*Pan troglodytes*; A), 29655 (*Pan paniscus*; B), and modern human male of European descent (*Homo sapiens*; C). Predicted thickness values for the composite skull of *Australopithecus africanus* (Sts 5 and Sts 52; D) are also shown. See abbreviations in Table 1.

## Discussion

Using Stephan’s (16) previously published soft tissue thicknesses for humans, a comparison can be made for 23 cephalometric points between *P. troglodytes* and *H. sapiens* means. Fig 5A shows a number of human thicknesses that are more similar to chimpanzees than reported in previous research measuring chimpanzee tissue thicknesses via ultrasound (17). The thickness of soft tissue in the area of the cheeks, which corresponds to landmarks ectomolare^2^ and ectomolare_2_, has been reported to differ between humans and chimpanzees with humans having thicker cheeks as a results of increased adipose tissue at this region (6, 17). However, our data show that mean thicknesses at ectomolare_2_ are identical between humans and chimpanzees and that ectomolare^2^ is only marginally larger in humans. Similarly, zygion is identical between species and not smaller in chimpanzees as previously reported. Of the larger differences shown in Fig 5B, there are only slight differences between human and chimpanzee means (minimum = 2 mm; maximum 7.8 mm). It is important to note that this comparison includes only 23 out of the 39 points that were measured in this study and, therefore, a more thorough comparison composed of a larger sample of human values may reveal less similarities than what is reported here. It may be argued also that, based on the similarities identified here, human and chimpanzee means are largely interchangeable and that this may appear like a valid option in the facial approximations of extinct hominids. However, we would like to remind the reader of two problems inherent in using means: 1) means have only been verified for a limited number of landmarks and therefore other regions of the face and head will need to be intuited or the thicknesses interpolated from species specific means; and, more importantly 2), means completely ignore variation among individuals. If we had interpolated chimpanzee means into our facial approximation of S9655 the average difference between the observed and predicted soft tissue thicknesses would have been 3.1 mm (minimum = 0.2 mm; maximum = 11.3 mm), which is higher than the average difference of 1.5 mm (minimum = 0 mm; maximum = 4.1 mm) that was produced using the regression models. Furthermore, with regard to our comparison of soft tissue thicknesses and craniometric dimensions between male and female chimpanzees, we have shown that in general they do not display sexual dimorphism and that this does not justify producing separate soft tissue prediction models for males and females because variation between sexes is negligible. In this respect, sexual dimorphism in chimpanzees is similar to modern humans (47), which corresponds with previous analyses of craniofacial sexual dimorphism among extant hominids (48, 49).

**Fig 5.**
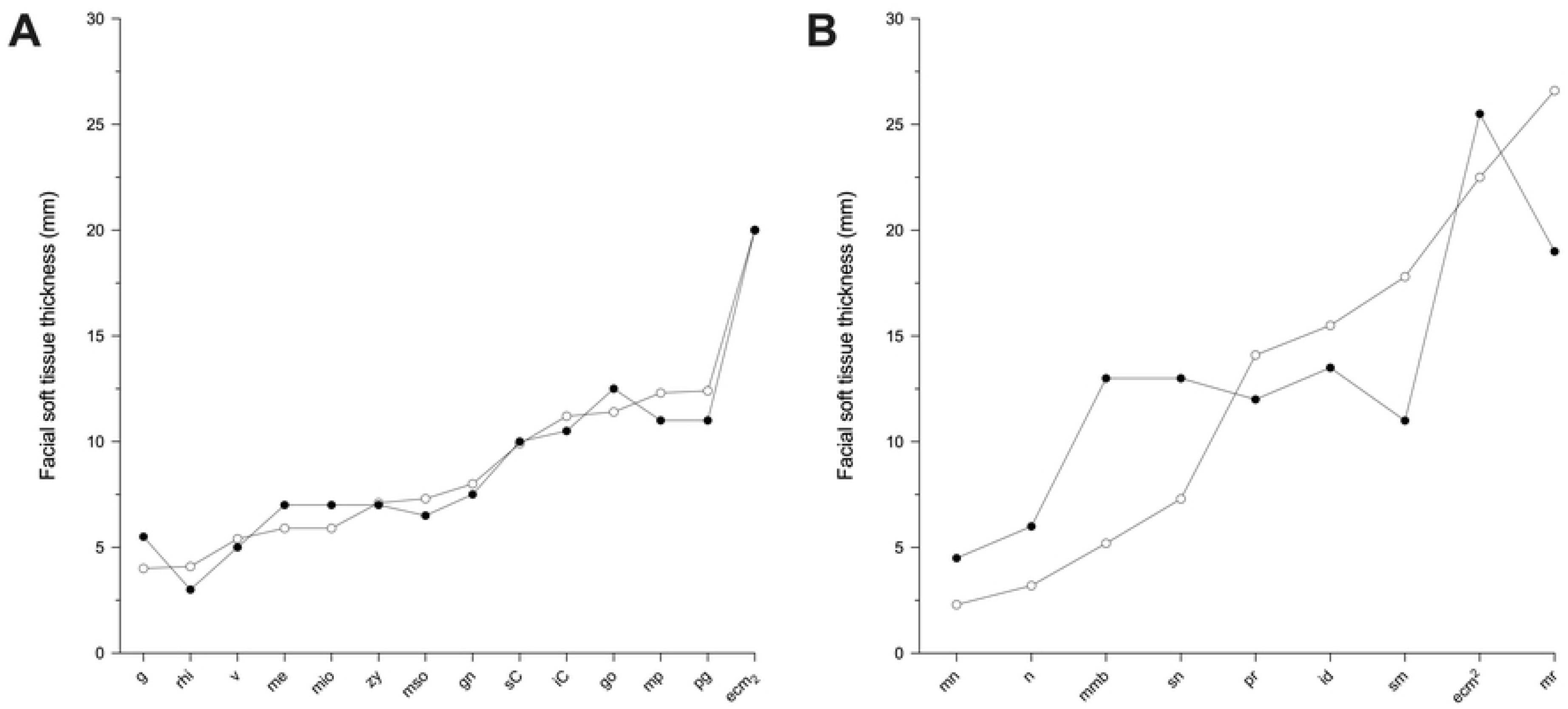
Line charts of *Pan troglodytes* (○) and *Homo sapiens* (●) means comparing values with differences < 2 mm (A) and differences > 2 mm (B).

The results of the out-of-group tests on the bonobo and human subjects suggest that chimp-derived regression models are compatible with species that have craniometrics that are more similar to chimpanzees than to those of modern humans. As is presented in Table 6 and Fig 6, chimpanzees, bonobos, and modern humans do not equally display disparate craniometric differences. Of the 15 craniometrics, S9655, had four outside the range of variation observed in this study sample of chimpanzees, whereas the human subject presented 11 craniometrics that were outside the range. The slight differences observed in the craniometric dimensions for S9655, however, did not appear to largely affect the predictive accuracy of the regression models, whereas large differences observed in the craniometric dimensions of the human subject produced large estimation errors. With the understanding that the craniofacial morphology of bonobos is similar to chimpanzees, particularly in cranial dimensions and the morphology of the masticatory apparatus, this result is to be expected. We admit that this test was conducted on two individuals only and that this may be perceived as weak evidence for regression model interspecies compatibility, however, we think it unreasonable to assert that all 26 regression models have performed fittingly on the bonobo subject and poorly on the human subject as a result of random chance.

**Fig 6.**
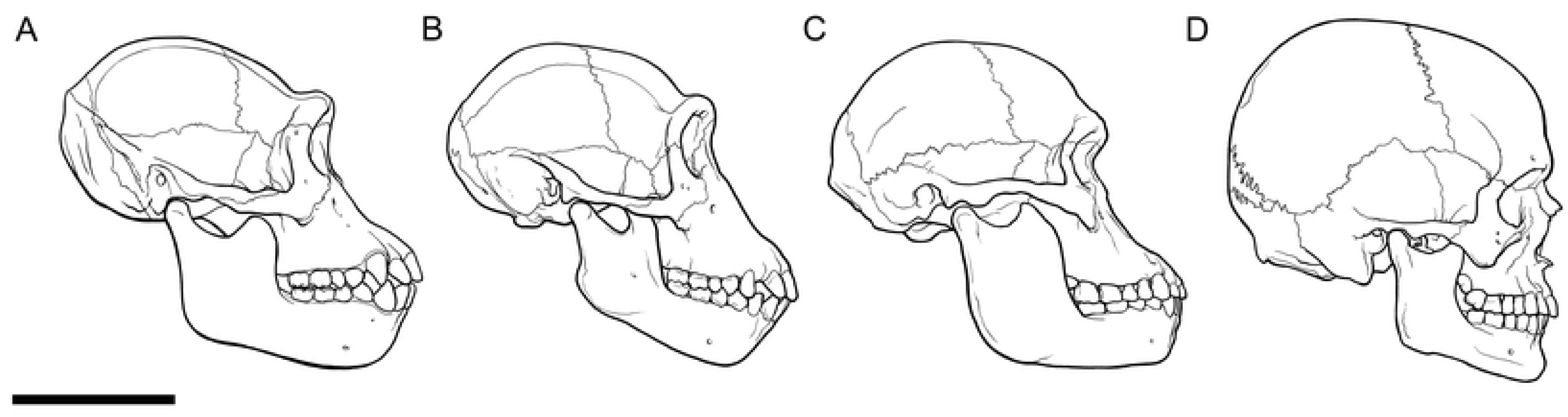
Skulls of PRI-Cleo (*Pan troglodytes*; A), 29655 (*Pan paniscus*; B), the composite skull of *Australopithecus africanus* (Sts 5 and Sts 52; C), and modern human male of European decent (*Homo sapiens*; D) in norma lateralis. Note the similarities and differences in the profiles of the facial projection and their implications for the thicknesses of the muscles that act on the masticatory system between these hominid species. Scale bar = 10 cm.

**Table 6.** Craniometrics taken from skulls of PRI-Cleo (Pan troglodytes), 29655 (Pan paniscus), composite skull of Australopithecus africanus (Sts 5 and Sts 52) and modern human male of European descent (Homo sapiens).

| Variable<sup>a</sup> | PRI-Cleo (*P. troglodytes*) | 29655 (*P. paniscus*) | Sts5/Sts52a (*A. africanus*) | Human subject (*H. sapiens*) |
| --- | --- | --- | --- | --- |
| zy-zy | 116.50 | **114.90** | 125.68 | 126.40 |
| g-n | 8.10 | 6.22 | 14.45 | 8.12 |
| v-gn | 178.80 | 191.55 | 195.55 | **224.30** |
| g-gn | 128.90 | 130.60 | 118.25 | **116.80** |
| v-po | 61.40 | 63.05 | 72.01 | **116.80** |
| v-sn | 132.80 | 141.50 | 132.90 | **174.40** |
| id-gn | 31.00 | **30.05** | **22.43** | **25.50** |
| al-al | 24.40 | **32.80** | 27.10 | **21.00** |
| v-g | 90.30 | 91.20 | 90.38 | **134.70** |
| c-c | 50.60 | 52.70 | **44.50** | **34.00** |
| v-n | 97.20 | 92.45 | 92.08 | **136.50** |
| go-go | 79.50 | 79.45 | 78.80 | 91.00 |
| n-pr | 77.60 | 86.60 | 79.85 | **67.20** |
| ft-ft | 51.60 | **72.05** | **63.98** | **93.50** |
| n-sn | 58.40 | 59.25 | **44.09** | 47.80 |

<sup>a</sup> See variable abbreviations in Table 1.
Bolded text indicates were craniometrics were outside ± two standard deviations from chimpanzee sample means.

Given that covariation between soft tissue thicknesses and craniometric measurements has been observed in both extant *Homo* and *Pan* species, we hold that it is reasonable to assume that such covariation was present in archaic hominids, such as Sts 5. We submit also that skull morphology is the prime determinant of regression model interspecies compatibility and that chimpanzee-derived regression models are valid for reconstructing the facial appearance of Sts 5. The justification for this is as follows. First, Sts 5’s craniometrics were just as different from chimpanzees as S9655’s were (Table 6; Fig 6). We must therefore agree with previous authors (50–52) in that *Pan* appears to be the most suitable extant hominid upon which extrapolations of covariation can be made for *A. africanus*. It is supported by the fact that since the chimpanzee-bonobo split *c*.2 Ma ago there have been no musculoskeletal changes in bonobos (23). If bonobos had gone extinct *c*.2 Ma ago, chimpanzee-derived regression models would still have produced an accurate result. Based on this and the closer affinity of the Sts 5 skull to *Pan*, any estimation errors in soft tissue thickness for Sts 5 are likely to be similar to or only slightly larger than those of S9655. Second, under the assumption that sexual dimorphism of soft tissue thicknesses in *A. africanus* did not differ significantly from chimpanzees and modern humans, it is clear that the lack of consensus surrounding the sex of Sts 5 did not affect the precision of the regression models’ predictions. Although we agree with Montagu (53) that the true soft tissue thicknesses for extinct hominids are largely unknowable, we argue that this fact does not diminish the utility of chimpanzee-derived regression models in formulating an informed hypothesis about the facial appearance of Sts 5 and other hominids with similar craniometrics.

To the best of the authors’ knowledge, regression models have not been used in the facial approximations of Plio-Pleistocene hominids prior to the present study. Earlier reconstructions have relied heavily on species specific means and/or comparative anatomy of primate muscle morphology (1). Here we will focus only on the latter as the limitations of means were discussed previously. The location and shape of muscle attachment areas on great ape skulls has been described in detail for bonobos (23), chimpanzees (25), orangutans (54), and gorillas (55) as well as in comparative anatomy textbooks (56). In any hominid facial approximation, there is an obvious importance in knowing the origin and insertion of the various muscles of the face and head between great apes and humans. However, knowledge of correct anatomical positions of individual muscles is not a substitute for specific estimates for the volumes of the muscles themselves and their coverings, namely the thicknesses of subcutaneous adipose tissues and epithelial linings. Gurche (1) reports being able to systematically determine the size, location and shape of muscles based on macroscopic surface markings on fossil bones. We will not be surprised if some readers support Gurche’s method as found throughout the facial approximation literature is the view that that the face can be reliably approximated from the construction of the facial musculature alone (13, 14, 30). However, as Ullrich and Stephan (57) have shown, this is a gross misinterpretation of the facial approximation method. In actual fact, facial approximation has always relied heavily on empirical data on soft tissue thicknesses (11, 12, 58). Gerasimov, for example, implemented soft tissue thickness measurements, only ever placed four muscles onto the skull (the masseter and temporalis muscles), and considered adding any further muscles, such as those of facial expression, to be pointless since their attachments to the skull are not visible. Furthermore, research attempting to recover the size and location of 92 muscles in human material reported that only 23 could be reliably reconstructed from bone alone (59). Gurche’s reconstructions are not necessarily illogical by any means but his approximations are not produced from direct observations of bone as commonly believed. While a practitioner’s sculptural skills and anatomical expertise is an obvious benefit in any facial approximation, in isolation the intuited use of this knowledge alone is highly vulnerable to subjective interpretation. For example, soft tissue may be added or subtracted based on personal preference. In contrast, the regression models of the present study provide direct evidence for the approximation of hominid soft tissues. These models may also be useful for studies reconstructing the physiology of extinct hominids. The masticatory system of Australopithecus, for example, may be analyzed in more detail by assigning empirical values to individual muscles of the head. Regression models for mid-ramus, temporal fossa and euryon reflect the volume of the masseter and temporalis muscles, which may be used to further analyze the biting performance of these hominid species. Regression models may also be extended in future studies to include the postcranial skeleton and improve upon current body mass estimates for extinct hominids (60).

The strong correlations observed in this study certainly raise questions about the claim that soft tissue thicknesses do not covary sufficiently enough with craniometric dimensions to improve soft tissue estimates in craniofacial identification (21). Given that correlation coefficients generated from regression analysis are sensitive to measurement error and that these errors can only detract from the strengths of association, it is possible that measurement error accounts for some, if not all, of the differences between the correlation coefficients of the present study and those reported in previous studies. However, this is difficult to evaluate until further analyses are made using human material and more reliable measurement methods.

It is important for us to be transparent about the limitation of the regression models. The aim of any facial approximation is to provide an accurate model of a complete subject. However, our regression models provide very little information about the facial features of hominids as they only provide a 3D silhouette upon which facial features can be built. In our approximation of subjects PRI-Cleo and S9655, the facial features can easily be extrapolated from photographic evidence of great apes as shown in the completed approximations in Fig 7A and B respectively. For Sts 5, however, practitioners of facial approximation have no direct information. Numerous facial approximation studies have developed methods for approximating the facial features in modern humans, however, the validity of these methods applied to other hominids has never been tested. Published methods include the approximation of eyeball diameter and anatomical placement in the orbits (61); eyebrow size, position and shape (62–65); nasal profile (12, 30, 66–70); mouth width and shape (11, 13, 30, 57, 71–73); and size and shape of the external ear (10-12, 65, 74, 75). Given that facial features are needed to complete any facial approximation, the interspecies compatibility of these methods is worthy of detailed examination in the near future to allow for complete approximations of Plio-Pleistocene hominids to be produced. Until then, the facial features presented in the completed facial approximation of Sts 5 shown in Fig 7C must obviously remain tentative.

**Fig 7.**
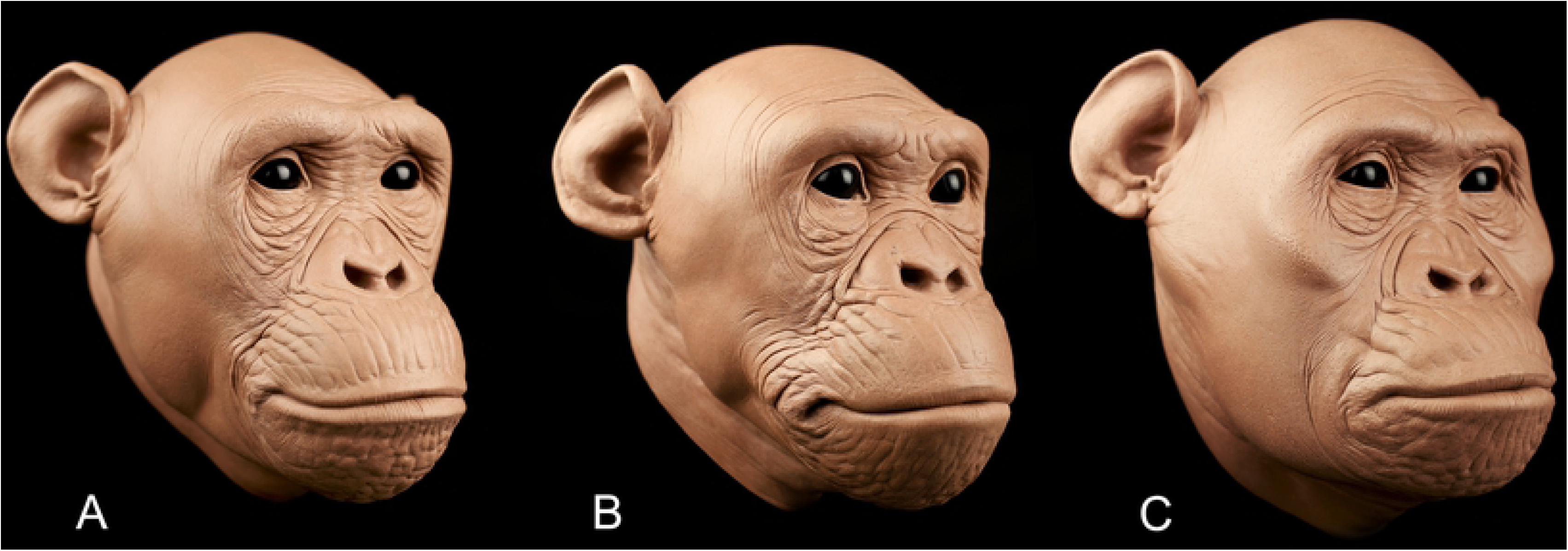
Completed facial approximations of PRI-Cleo (*Pan troglodytes*; A), 29655 (*Pan paniscus*; B), and composite skull of *Australopithecus africanus* (Sts 5 and Sts 52; C) with facial features extrapolated from photographic evidence of chimpanzees and bonobos. Please note that the facial features of *A. africanus* are highly approximate. Scale bar = 10 cm.

Second, craniofacial morphology among Plio-Pleistocene hominid taxa is highly variable and as such not all fossil craniometrics may fall inside or close to the range modelled in the present study’s regression analysis. Those hominid crania with craniometrics that are at or outside of the extreme ends of the independent variable are likely to produce large estimation errors in soft tissue thickness. As mentioned previously, the craniometrics of Sts 5, as well as subject S9655, are within and close to the range of variation observed in the present study sample of chimpanzees. However, this was not the case for the modern human, which resulted in poor approximation of soft tissue thicknesses, and will not be the case for all hominid skulls, especially specimens with craniofacial morphologies like those exhibited in *Paranthropus boisei* (76, 77). Repeating stepwise multivariate regression analyses on other extant apes with craniometrics that approximate these fossils, such as gorillas and orangutans, is one possible solution to be explored in future studies.

Finally, it cannot be assumed that correlations observed in adult subjects scale isometrically for very young hominid skulls like that of the Taung fossil or DIK-1 (78, 79). Changes in modern human soft tissue thicknesses between 0 and 19 years of age have been shown to be small and relatively constant throughout ontogeny (16, 80), but their relations to substantially growing craniometric dimensions may not be the same as in adults. Similarly, thickness changes throughout ontogeny in other primates are unknown, therefore the regression models of the present study may only be viable for approximating adult hominid faces.

## Conclusions

The results of this study show that soft tissue and craniometric measurements covary in chimpanzees, which confirms that such covariation is uniformly present in both extant *Homo* and *Pan* species. Chimpanzee-derived regression models appear to be compatible with bonobos but show a marked decrease in predictive accuracy in humans, suggesting that regression model reliability is dependent on craniometric similarity. As the craniometrics dimensions of early hominids, such as South African australopithecines, are more similar to chimpanzees than those of humans, chimpanzee-derived regression models may be used to approximate their craniofacial anatomy. Additional relationships between soft tissue thickness and craniometric dimensions of non-human hominids may contribute to more precise facial approximations of early hominids bringing them even closer to standards of objectivity used in forensic sciences. It is hoped that the results of the present study and the reference dataset for facial soft tissue thicknesses of chimpanzees it provides will encourage further research into this topic.

## Author contributions

All authors contributed to the conception and design of the project, interpreted research data, edited the manuscript and approved the final version of the manuscript. R.C. drafted the manuscript, collected measurement data and performed statistical analyses of these data being advised by M.H. G.V. prepared skulls for 3D printing and performed the 3D reconstructions presented in Figures 3 and 7.

## Competing interests

The authors declare they have no competing interests.

## Acknowledgments

We would like to thank the Kyoto University Primate Research Institute (Kyoto, Japan), Morphosource (https://www.morphosouce.org) and Stefano Benazzi (University of Bologna) for facilitating the acquisition of digital data. We would like to acknowledge Ian Linke (University of Adelaide) for providing access to the 3D printing lab, Victor Surovec (Arizona State University) for providing access to the Makerspace, Dan Collins (Arizona State University) for providing access to the Vizproto Lab, and Michal Dutkiewicz for the illustration presented in Figures 1 and 5. Aspects of this work were funded by the University of Adelaide.

